# Intra-subgenome regulation induces unbalanced expression and function among bread wheat homoeologs

**DOI:** 10.1101/2024.08.01.606124

**Authors:** Xiaoming Wang, Yuxiu Liu, Peng Zhao, Wenyang Hou, Mingzhu Cheng, Xue Shi, James Simmonds, Philippa Borrill, Wanquan Ji, Shengbao Xu

## Abstract

The differential expression of homoeologous genes confers expression plasticity and facilitates the adaptation and domestication of major polyploid crops. However, how this homoeolog expression bias (HEB) is regulated remains elusive. Here, transcriptome analysis of 406 wheat (*Triticum aestivum*; AABBDD) accessions revealed great variation in HEB among accessions. We identified 14,727 QTLs regulating HEB (hebQTLs), indicating that HEB is genetically regulated and can be predicted using genotyping data. The hebQTLs only regulate the expression of homoeologs in the same subgenome and downregulate their expression to result in HEB, suggesting that intra-subgenomic rather than inter-subgenomic interactions induce HEB. Furthermore, non-hebQTL-regulated homoeologs have stronger biological functions, are under higher selection pressure and exhibit lower genetic diversity than hebQTL-regulated homoeologs and compensate for the downregulated expressions of hebQTL-regulated homoeologs. Our findings reveal how homoeolog expression is coordinated at the genetic level and provide an avenue for leveraging HEB to improve polyploid crops.

## Main

Polyploid crops, including wheat (*Triticum aestivum*), cotton (*Gossypium hirsutum*), potato (*Solanum tuberosum*), oilseed rape, tobacco, sugarcane, apple, banana and others, play vital roles in feeding the world’s population. The polyploid genome contains multiple gene copies known as homoeologs. This provides redundant genetic pools and improves the expression flexibility of genes due to the differential expression levels of homoeologs^1–8^. These attributes are thought to confer adaptive plasticity and fitness to crops in a new environment, which is critical for the broader geographic distribution of polyploid crops compared to their diploid relatives^2,9–14^. An in-depth understanding of the expression flexibility of homoeologs could help leverage the advantages of polyploidy to improve crop production and help crops adapt to climate change.

Bread wheat (*Triticum aestivum* ssp. *aestivum*, 2*n* = 6*x* = 42,AABBDD), the most widely cultivated crop on Earth, contributing approximately one-fifth of the total calories consumed by humans, is an allohexaploid species that originated from two hybridization events^15^. The first hybridization, between *Triticum urartu* (AA) and a close relative of *Aegilops speltoides* (BB), gave rise to tetraploid wheat (AABB), and the second hybridization between tetraploid wheat and *Aegilops tauschii* (DD) generated allohexaploid wheat^6,16,17^. Most wheat genes are present in three copies (homoeologs), constituting a triad. The available whole genome sequence^15^, along with the high degree of autonomy in the subgenomes^17^ and more recent polyploidization^18^, make wheat an informative system for studying the effects of polyploidy on gene expression.

We previously observed homoeolog expression bias (HEB) in ∼30% of wheat triads, meaning that one homoeolog has a higher or lower expression level than the other two homoeologs^11^. We also determined that the types of HEB for some triads vary among 15 wheat tissues^11^: for example, the expression of a homoeolog in subgenome A was suppressed in roots, while the expression of homoeologs in both subgenomes A and B was suppressed in leaves, leaving the HEB regulation as an intriguing question. Besides, it was reported that HEB is involved in regulating agricultural traits and stress responses^18–26^, including the adaptation of wheat flowering to a range of environmental conditions, a process regulated by *VRN1* and *PPD1*^27,28^. Therefore, the factors regulating HEB are also important for understanding and leveraging the broader adaptation and agricultural trait variations of wheat. Most studies of HEB regulation have shown that transposable elements and epigenetic factors are related to HEB^11,24,29–32^. However, whether, to what extent or how HEB is regulated by genetic factors is largely unknown. A recent study showed that cis-subgenomic SNPs explained a higher proportion of gene expression variance than inter-subgenomic SNPs and suggested that the discordant expression of homoeolog pairs is associated with the accumulation of cis-rather than trans-regulatory variants in wheat^33^, providing initial clues about the genetic regulation behind HEB.

Here, based on our previous root transcriptomes of 406 worldwide wheat accessions^34^, the types of HEB were found to vary strongly among accessions. Next, we developed a model to transfer the extent of HEB among accessions to a one-dimensional value. A genome-wide association study (GWAS) revealed 14,727 quantitative trait loci regulating HEB (hebQTLs), shedding light on the genetic factors regulating HEB. Further analysis showed that these hebQTLs mainly regulate the expression variance of homoeologs located in the same subgenome rather than the other two homoeolog copies, suggesting that the driving force of HEB is intra-rather than inter-subgenome interactions. Finally, we provide evidence that HEB and hebQTLs are associated with root-related traits and that the hebQTL-regulated homoeolog in a triad is under relaxed selection pressure and accumulates more sequence variants than the other two homoeolog copies, thereby having less significant effects on root-related traits. Our results provide an understanding of the genetic regulation of HEB among accessions, laying the foundation for improvement and adaptive fitness of polyploid crops via the genetic manipulation of HEB.

## Results

### The three wheat subgenomes play unbalanced roles in root development

Analysis of the root transcriptomes of 406 worldwide wheat accessions produced in our previous study^34^ revealed that 68,418 (65.04%) high-confidence genes were expressed in roots based on transcripts per million (TPM) >0.5 in at least 5% of accessions, including 22,885 (64.75%), 22,936 (64.35%) and 22,597 (66.05%) genes located in the A, B and D subgenome, respectively (http://resource.iwheat.net/WGPD/). The mean expression levels of genes in subgenome D were slightly higher than that for either subgenome A or B (Fig. 1a). Notably, for the top 10% most highly expressed genes in each subgenome, 12.84% (in subgenome D) to 14.83% (in subgenome A) of genes lacked homoeologous copies in the highly expressed gene set of either of the two other subgenomes (Fig. 1b,c and Table S1). These subgenome-private highly expressed genes were mainly enriched for the metabolism-related GO terms (Fig. S1a and Table S2).

**Figure 1.**
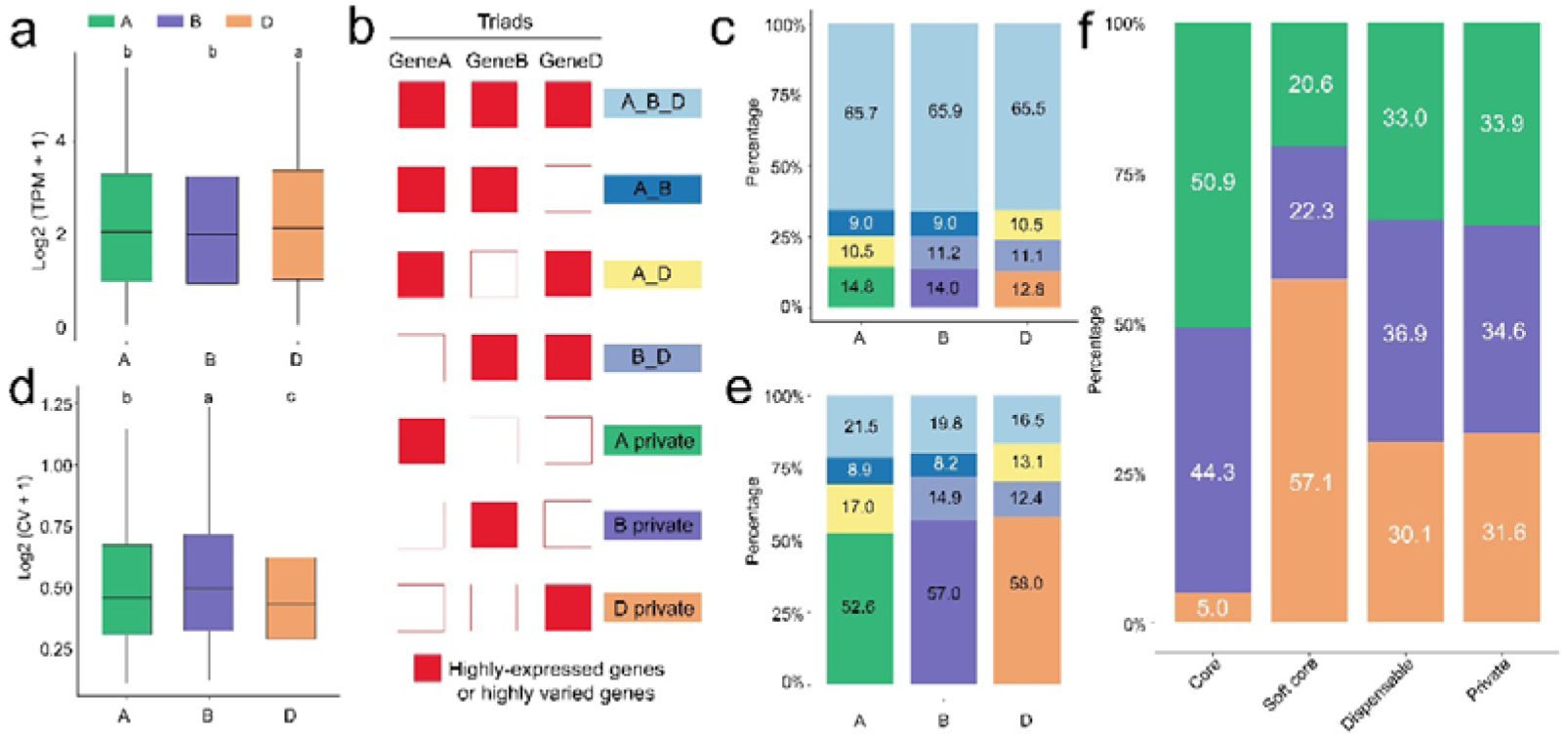
Imbalance of the three wheat subgenomes based on root transcriptome data. **a**, Comparison of gene expression levels among the three wheat subgenomes. **b**, The model deciphering the categories of homoeologous copies between the top 10% most highly expressed or highly varied genes in the three subgenomes. **c**, The percentages of the top 10% most highly expressed genes which have or have no homoeologous copies in the highly expressed gene sets of the two other subgenomes. **d**, Comparison of the coefficient of variation of gene expression levels among the three subgenomes. **e**, The percentages of the top 10% most highly varied genes which have or have no homoeologous copies in gene sets of the two other subgenomes. **f**, Distributions of the four gene categories in pan-transcriptome analysis among the three subgenomes. In **a** and **d**, different lower-case letters indicate significant differences (*P* < 0.05) based on least significant difference (LSD) test for multiple comparisons. Boxes show median values and the interquartile range. Whiskers show minimum and maximum values, excluding outliers.

The genes in subgenome B showed significantly higher expression variance among the accessions than genes in subgenome A or D (Fig. 1d), consistent with the known higher genetic diversity of subgenome B^16^. Among the top 10% of genes with the highest expression variance in each subgenome, 52.56%, 57.01% and 58.00% of those in subgenomes A, B and D, respectively, lacked homoeologous copies in gene sets in any of the other two subgenomes (Fig. 1b,e and Table S3). The enriched GO terms of these subgenome-private genes included "stress response", "regulation of cell death", "salicylic acid biosynthetic process", "hydrogen peroxide catabolic process", and "nutrient reservoir activity" (Fig. S1b and Table S4).

We investigated the expression breadth of each gene and defined 19,592, 25,759, 23,067 and 36,783 genes as core, soft core, dispensable and private genes, indicating that the genes are expressed in all, ≥90% and <100%, ≥5% and <90%, and <5% of the accessions examined, respectively (Table S5). The proportions of core genes in subgenomes A and B were comparable but much larger than that in subgenome D (Fig. 1f), perhaps because subgenome D was the most recent subgenome to be incorporated into the wheat genome^6,16,17^. The private genes were present in similar proportions in the three subgenomes. Although some similar enriched GO terms were identified, the private genes were also enriched in different GO terms, including “regulation of cell morphogenesis”, “auxin biosynthetic process” and "salicylic acid metabolic process" for subgenome A; “response to salicylic acid”, "electron transfer" and "protein catabolic process" for subgenome B; and “response to ethylene”, "phosphate ion binding" and "translation" for subgenome D (Fig. S1c and Table S6). These results point to unbalanced roles for the three subgenomes in root development.

### HEB categories vary among wheat accessions

To further investigate the imbalance of the three subgenomes, we filtered the reported triads (1:1:1 correspondence across the three homoeologous subgenomes)^11^, retaining 14,739 triads with a summed expression of >0.5 TPM across the triad (Table S7). We also standardized the relative expression levels of the A, B and D homoeologs so that the sum was 1.0 in each accession, making homoeolog expression levels comparable across triads and accessions. The 14,739 triads were divided into seven categories by calculating the mean expression level of each homoeolog across the accessions and determining the triad’s position in the ternary plot, as previously described^11^, including 10,931 balanced triads, 940 homoeolog-dominant triads (307, 279 and 354 for A-, B- and D-homoeolog dominant, respectively) and 2,868 homoeolog-suppressed triads (1044, 1181 and 643 for A-, B- and D-homoeolog suppressed, respectively) (Fig. 2a and Table S7). Consistent with our previous findings about HEB among 15 wheat tissues^11^, the genes from balanced triads were expressed across a broader range of accessions and had higher absolute expression values, but lower expression variance among accessions, compared to the genes from unbalanced triads (Fig. S2). Likewise, analysis of absolute expression (TPM) shows that HEB resulted from relatively low expression of one or two homoeologs instead of expression increases (Fig. S2b). We refer to the homoeologs with lower expression in homoeolog-suppressed and homoeolog-dominant triads as suppressed-homoeologs and to those with higher expression within the triads as non-suppressed-homoeologs hereafter (Fig. 2b).

**Figure 2.**
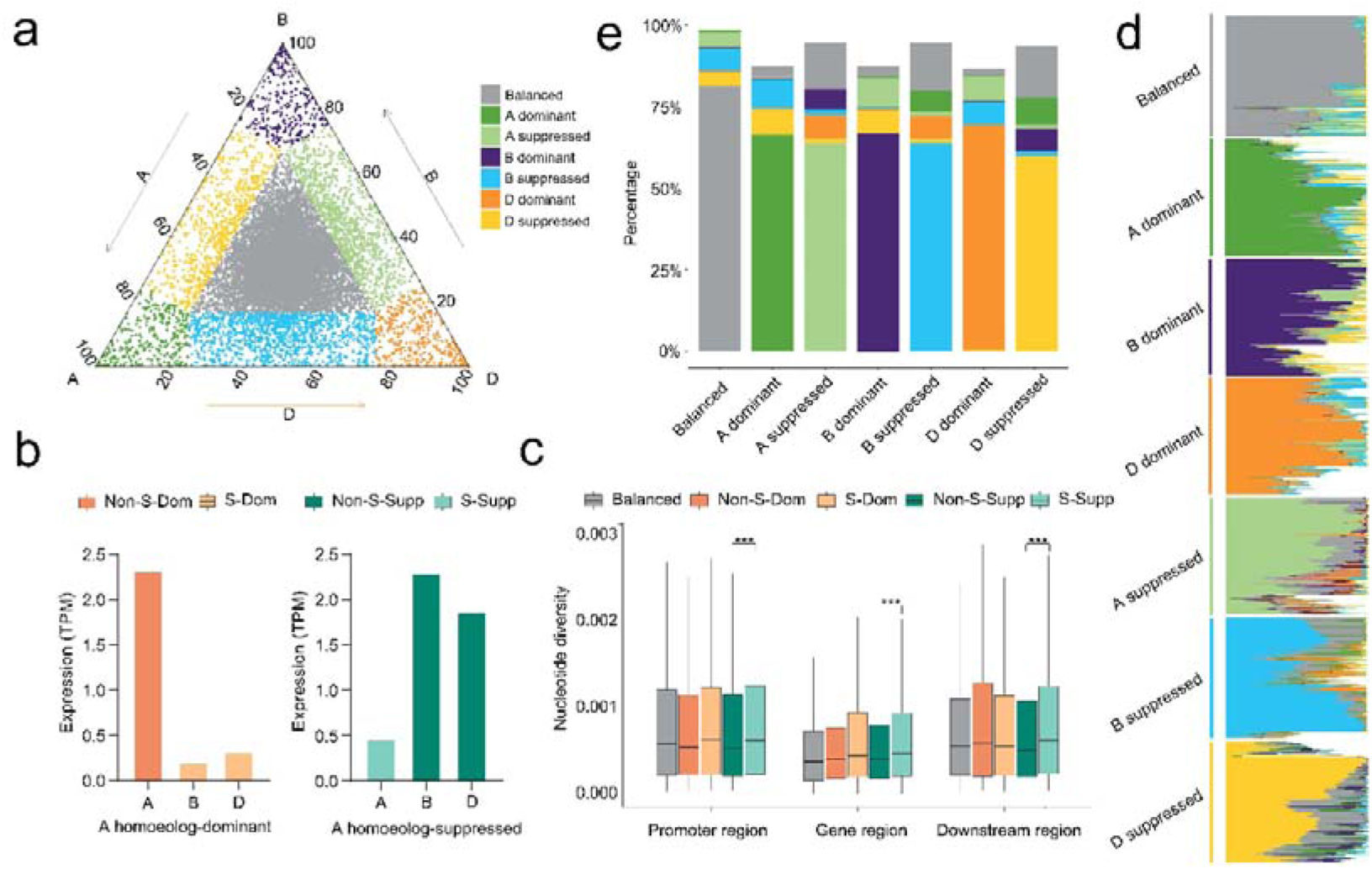
Homoeolog expression bias (HEB) varies among wheat accessions. **a**, Seven triad categories defined with the triad’s positions on the ternary plot. Each point represents a triad, corresponding to the relative expression abundance of A, B and D homoeologs. **b**, The model deciphering the definitions of "S-Dom", "Non-S-Dom", "S-Supp", and "Non-S-Supp" in the homoeolog-dominant and homoeolog-suppressed triads, taking an A-homoeolog-dominant triad and an A-homoeolog-suppressed triad as examples. **c**, Comparisons of genetic diversity between homoeologs with suppressed vs. non-suppressed expression from different triad categories. Asterisks indicate significant differences: \*\*\**P* < 0.001 based on Student’s *t*-test. Boxes show median values and the interquartile range. Whiskers show minimum and maximum values, excluding outliers. **d**, Conservation of global triad category (labelled at left) among the 406 wheat accessions. Each row represents a triad, and each column represents the triad category in the relative accession (accession category). The white cells in the heatmap represent an undefined triad category due to the sum expression (TPM) of the three homoeologs in the relative accession being <0.5. **e**, The percentages of accession categories in each of the seven global triad categories.

The release of new wheat resequencing data^16^ gave us the opportunity to investigate the reasons for the downregulated expressions of suppressed-homoeologs. We determined that in homoeolog-suppressed triads, genetic diversity was significantly higher in both the coding regions and the up- and down-stream regions of suppressed-homoeologs as compared to non-suppressed-homoeologs (Fig. 2b,c). For homoeolog-dominant triads, the same trend was observed in the coding and upstream regions, although the difference was not significant. Consistent with the higher genetic diversity of subgenome B^16^, the number of unbalanced triads containing suppressed B-homoeologs, A- and D-homoeolog-dominant and B-homoeolog-suppressed triads, was significantly larger than the number of unbalanced triads with other three categories (Chi-square test, *P*-value = 4.01e-10). These findings suggest a correlation between higher genetic diversity and lower homoeologs expression.

Next, we investigated whether and to which extent the HEB categories varied among accessions, which is fundamental for investigating the genetic regulation of HEB. We compared the triad category calculated using the global mean TPM vs. the 406 categories calculated using the TPM of each accession. With the criterion that at least 5% of the accessions belonged to a different triad category from that calculated using the global mean TPM, the HEB categories of 18.78%, 31.07%–33.65% and 36.16%–40.19% of the balanced, homoeolog-dominant and homoeolog-suppressed triads, respectively, were variable among accessions (Fig. 2c,d and Table S8). In detail, the unbalanced triads shifted most often to the three adjacent categories on the ternary plots, ranging from 19.13% to 22.25% for the homoeolog-dominant and 30.16% to 33.66% for the homoeolog-suppressed triads. In a few cases, the unbalanced triads shifted to the three non-adjacent categories, ranging from 1.19% to 2.13% for the homoeolog-dominant triads and 3.14% to 3.44% for the homoeolog-suppressed triads (Fig. 2d and Fig. S3). These data indicate that the triads’ HEB categories greatly varied across accessions, and the ratio was higher in unbalanced triads than that in balanced triads.

### Intra-subgenome regulation induces the unbalanced expression of the three homoeologs

To quantify the extent of HEB variation among accessions, we calculated the mean Euclidean distance (ED) between the triad’s position in each accession and its global average position on the ternary plot for each triad using our previously developed method^11^, generating a distribution of mean distances in which larger distances indicate higher variations (Fig. 3a). The triads with the 20% shortest and longest mean distances across accessions were defined as stable and dynamic triads, respectively (Table S9). As expected, stable triads were enriched for GO terms related to housekeeping functions, e.g., translation and metabolism, whereas dynamic triads were mainly enriched for environmental responses, cell population proliferation and hormone signalling (Fig. S4 and Table S10). Further analysis showed that genes from stable triads had much higher absolute expression levels than those from dynamic triads (Fig. 3b). Moreover, suppressed-homoeologs in dynamic triads showed significantly greater expression variation than non-suppressed homoeologs, suggesting that the expression variation of suppressed-homoeologs underlies changes in the HEB category (Fig. 3c).

**Figure 3.**
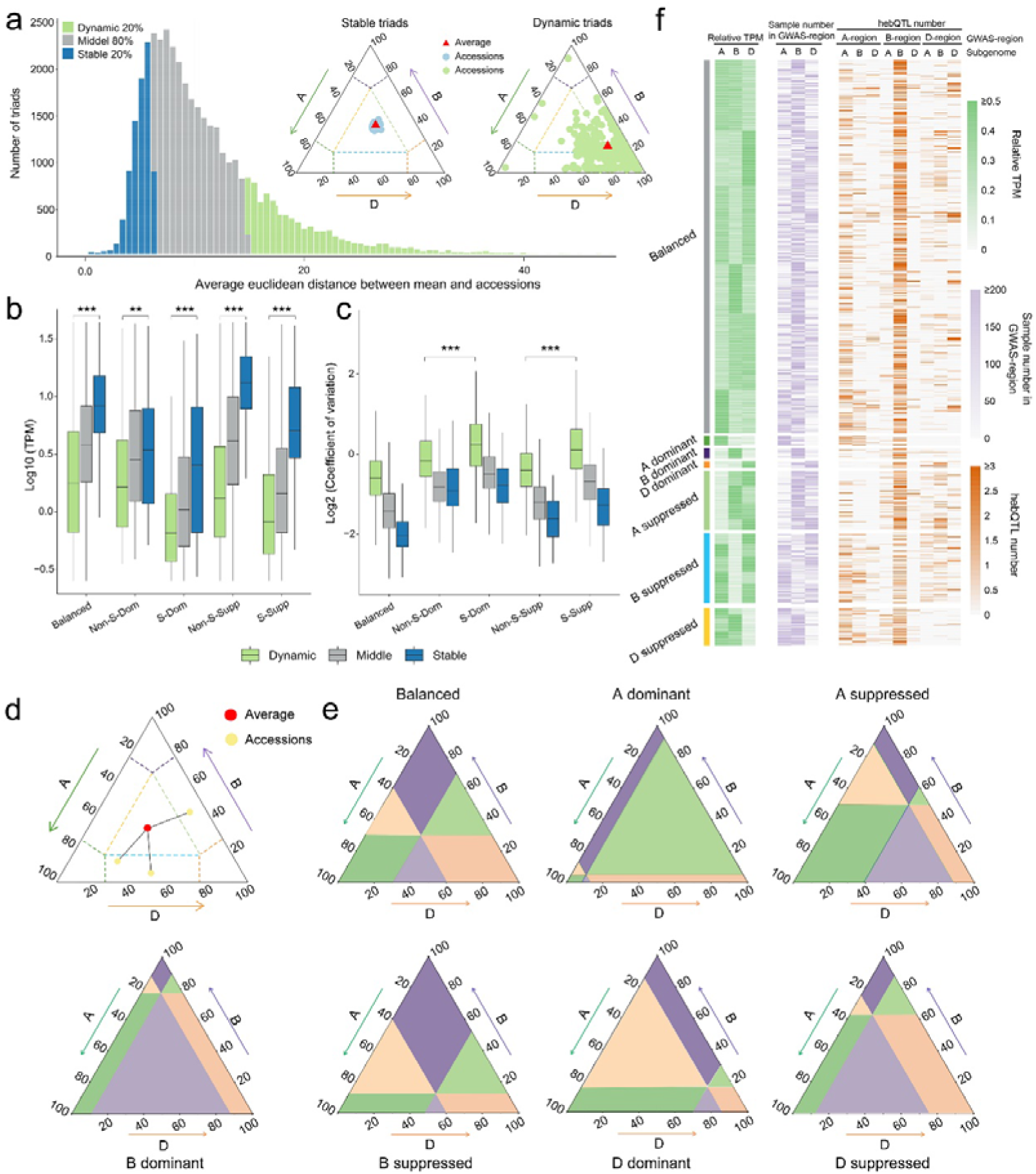
Identification of QTLs regulating HEB variation among accessions (hebQTL). **a**, Distribution of mean distance of triad variation across 406 accessions. The two ternary plots show a stable triad and a dynamic triad, respectively, in which "Mean" represents the global average point and "Accessions" represents the points corresponding to the 406 accessions. **b-c**, Comparisons of the absolute expression levels (**b**) and the coefficient of variation (**c**) of homoeologs with suppressed vs. non-suppressed expression from different triad categories. Definitions of "S-Dom", "Non-S-Dom", "S-Supp", and "Non-S-Supp" refer to Fig. 2b. Asterisks indicate significant differences: \*\**P* < 0.01, \*\*\**P* < 0.001 based on Student’s *t*-test. **d**, A model showing the direction attribute of Euclidean distance between the accession point and the global average point on the ternary plot. **e**, Models showing how the ternary plot was divided into six regions based on the global average point for seven triad categories. The two head-to-head regions (with similar colours) form a single GWAS region. **f**, Information of the identified hebQTLs, including which GWAS-regions they were identified and which subgenome they are located on. GWAS-region A, B and D represent the GWAS-region containing the vertex A, B and D of the ternary plot in **e**.

We then investigated the genetic factors regulating the HEB variation of each dynamic triad by considering the ED of each accession’s position from the global average position on the ternary plot as a phenotype and associating these phenotypes with our previously identified genetic variants^34^ by GWAS. However, in addition to the length values, the ED on the ternary plot also have direction attributes, which represent the HEB category in each accession (Fig. 3d). To address this issue, we divided the ternary plot into six regions based on the global average position of a given dynamic triad, making the ED of accession’s positions in each region have highly similar directions (Fig. 3e). We then considered two head-to-head regions as a single GWAS region and adjusted the ED values in these two regions as positive and negative values, respectively. Therefore, the ternary plot was divided into three GWAS regions (Fig. S5). Finally, we used the adjusted ED values in each GWAS region to perform GWAS and merged the significantly associated SNPs into QTLs via two steps (see Online Methods). Considering that these QTLs regulate the HEB variation among accessions, we will refer to them as hebQTLs hereafter. We identified 14,727 hebQTLs for 2,043 dynamic triads, including 10,261, 580 and 3,886 hebQTLs for 1,346 balanced, 96 homoeolog-dominant and 601 homoeolog-suppressed triads, respectively (Fig. 3f, Fig. S6 and Table S11). This is the first report, to our knowledge, of the genetic variants that regulate HEB variation.

The identification of hebQTLs provided us with the opportunity to investigate whether the driving force of HEB arises from intra-subgenome or inter-subgenome interactions. We estimated the genetic effects of each hebQTL on the expression variations of the three homoeologs within the relative triad using the general linear model and determined that a hebQTL only regulates the expression variation of the homoeolog from the same subgenome (Fig. 4a,b). Furthermore, we analysed the overlap between a triad’s hebQTLs and eQTLs (expression QTLs) that regulate the expression variations of the three homoeologs, finding that hebQTLs overlapped mainly with eQTLs of homoeologs in the same subgenome (Fig. S7). These results demonstrate that the driving forces of HEB arise from intra-subgenome interactions.

**Figure 4.**
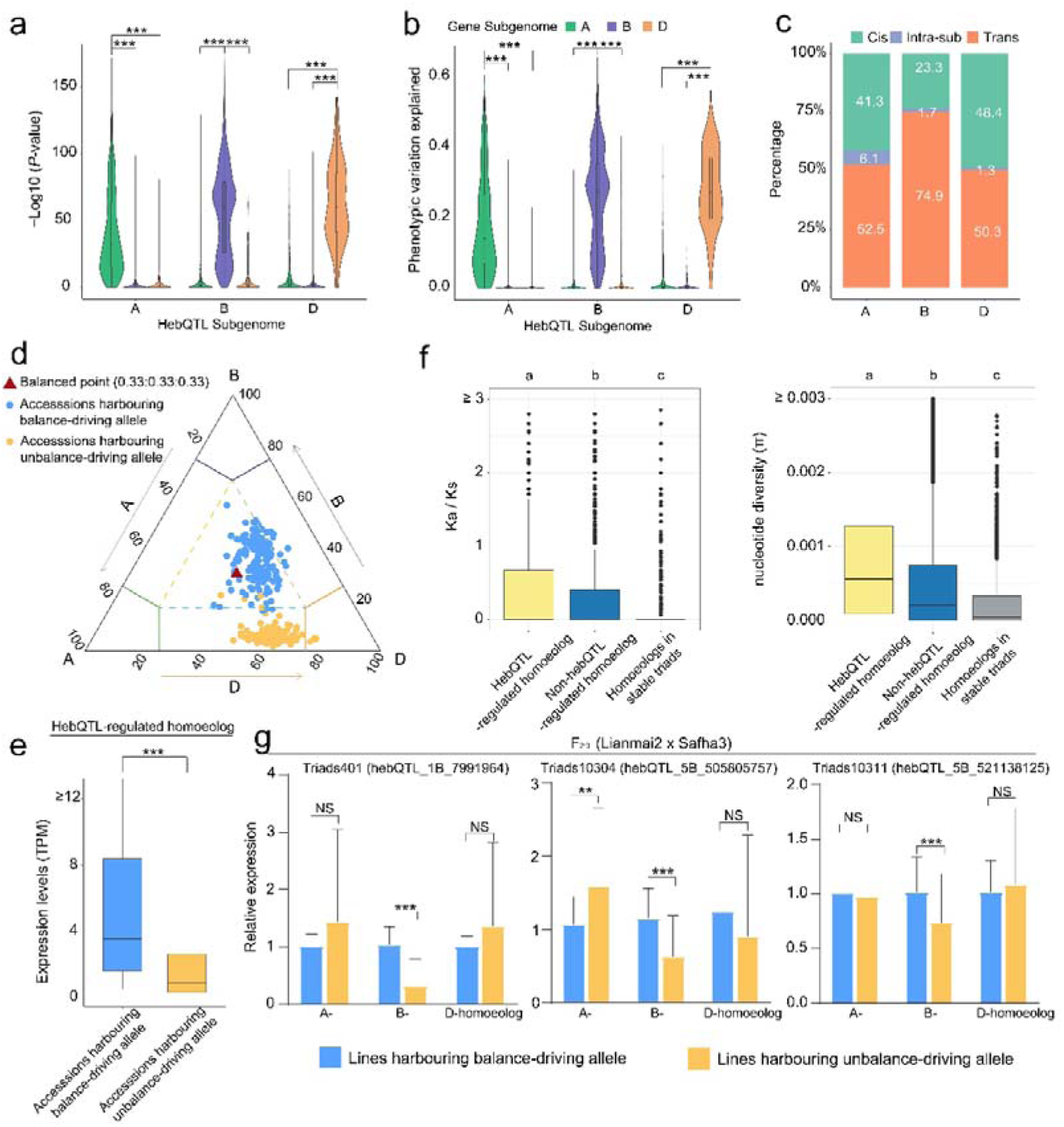
Characterization and validation of hebQTLs. **a**,**b**, *P*-values (**a**) and phenotypic variation explained values (PVE, **b**) of the genetic effects of the identified hepQTLs on expression variation of the three homoeologs among accessions. The genetic effects were estimated using the general linear model. **c**, Proportions of the three types of hepQTLs in the three wheat subgenomes. **d**, The model deciphering the definition of unbalance- and balance-driving alleles. **e**, Comparison of the expression levels of hepQTL-regulated homoeologs between accessions harbouring unbalance- and balance-driving alleles. **f**, Comparison of *K*_a_/*K*_s_ and nucleotide diversity between hepQTL-regulated homoeologs and non-hepQTL-regulated homoeologs (the two other homoeologs that were not regulated by a given hepQTL). The values of the three homoeologs from stable triads were used as a control. Different letters indicate significant differences (*P* < 0.05) based on least significant difference (LSD) test for multiple comparisons. **g**, Comparison of the expression levels of the three homoeologs between homozygous F_2:3_ lines carrying alternative hepQTL alleles. In **a**, **b**, **e** and **g**, asterisks indicate significant differences: \**P* < 0.05, \*\**P* < 0.01, \*\*\**P* < 0.001, NS, *P* > 0.05, based on Student’s *t*-test.

Given the hebQTLs only regulated the expressions of same-subgenome homoeologs, the identified hebQTLs were divided into three types: cis-hebQTLs (23.34%–48.47%) located ±5 Mb from the genes they regulate; trans-hebQTLs (50.25%–74.94%) located on the same chromosome as the genes they regulate; and intraSub-hebQTLs (1.28%–6.10%) located on different chromosomes but in the same subgenome as the genes they regulate (Fig. 4c and Table S11).

Next, we named the two alleles of the lead SNP for each hebQTL the unbalance-driving and balance-driving allele, respectively, meaning that accessions harbouring unbalance-driving and balance-driving alleles are located farther from and closer to the centre point of the ternary plot, respectively (Fig. 4d). The absolute expression levels of the hebQTL-regulated homoeologs were significantly lower in accessions harbouring the unbalance-driving allele than in those harbouring the balance-driving allele, suggesting that the hebQTL downregulates the homoeolog in the same subgenome to result in HEB (Fig. 4e). To investigate the reasons of hebQTL-regulated homoeologs were downregulated, we calculated the *K*_a_/*K*_s_ values of the three homoeologs, finding values of hebQTL-regulated homoeologs were higher than that of the other two homoeologs (Fig. 4f). Genetic diversity analysis revealed the same trend as for the *K*_a_/*K*_s_ values (Fig. 4f). These results suggest that the more relaxed selection pressure and more accumulated genetic variants generated the unbalance-driving alleles, which downregulated the hebQTL-regulated homoeologs and resulted in HEB, consistent with the former results that HEB resulted from relatively low expression of one or two homoeologs instead of expression increases.

To experimentally validate the downregulated effects of unbalance-driving alleles on the hebQTL-regulated homoeologs, we selected three hebQTLs from three dynamic triads. We constructed an F_2:3_ bi-parental segregation population by crossing Lianmai2 and Safha3, which harbour alternative alleles for the three hebQTLs. We measured the expression levels of the three homoeologs of each triad by qRT-PCR and compared them between homozygous F_2:3_ lines carrying alternative hebQTL alleles, respectively (Tables S12 and S13). The expression levels of the hebQTL-regulated homoeolog were significantly lower in lines harbouring the unbalance-driving vs. balance-driving allele (Fig. 4g). In contrast, the expression levels of the two other homoeologs which were not regulated by the hebQTL (non-hebQTL-regulated homoeologs), did not show significant difference with the exception of one homoeolog. Meanwhile, the positions of lines harbouring alternative alleles on the ternary plot were clearly separated (Fig. S8). These results support the notion that the unbalance-driving alleles of hebQTLs downregulate the expression of homoeologs in the same subgenome, resulting in HEB.

### hebQTL-regulated homoeologs have weaker effects in regulating root development than the two other homoeologs

To more deeply explore the contribution of HEB to adaptive plasticity and fitness in wheat^18–26^, we estimated the genetic effects of the identified hebQTLs on root-related traits by investigating whether the traits were significantly different between accessions harbouring alternative hebQTL alleles using the general linear model, which accounts for population structure, using our previously described method^35^. Of the hebQTLs identified, 55.37% had genetic effects on at least one root-related trait (FDR <0.05) (Table S14). We then calculated the significance of associations between the ED values in each GWAS-region and the root-related traits, finding 634 pairs of significant associations from 353 triads (Fig. S9 and Table S15). To experimentally validate these identified genetic effects and associations, we took the above-mentioned F_2:3_ segregation population and the three hebQTLs which were identified to have genetic effects and found that the homozygous lines harbouring alternative alleles showed significantly different root-related traits (Fig. 5a). Then, the ED values of the homozygous F_2:3_ lines on the ternary plot calculated with the relative expression levels of the three homoeologs (via qRT-PCR) were significantly associated with at least one root-related trait (Fig. 5b). These results provide direct evidence for the correlation between HEB and agricultural traits.

**Figure 5.**
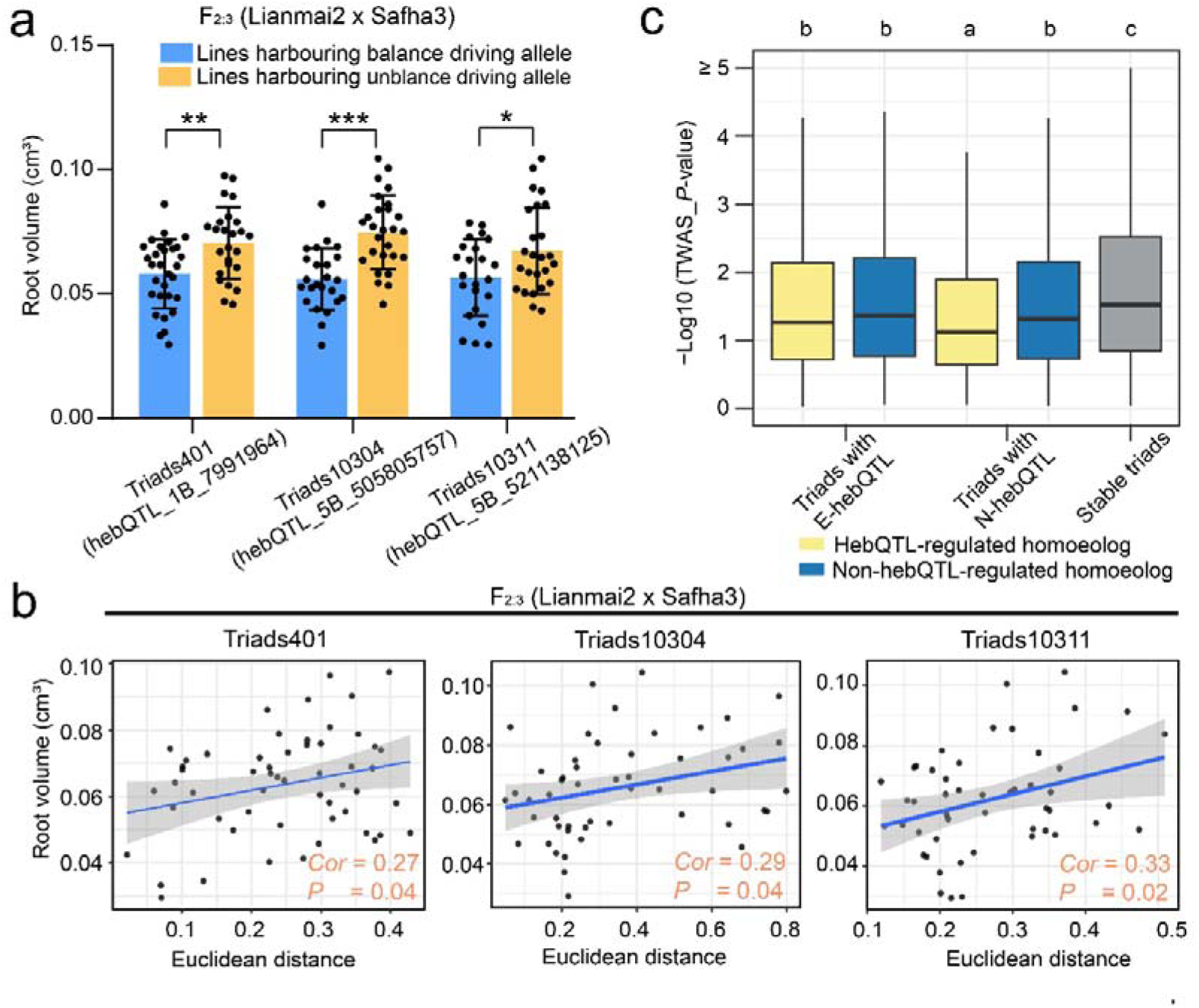
hepQTL-regulated homoeologs underlie the genetic effects of hepQTLs. **a**, Comparisons of root volume between homozygous F_2:3_ lines harbouring alternative alleles of the three hepQTLs. Asterisks indicate significant differences based on Student’s *t*-test (\**P* < 0.05; \*\**P* < 0.01; \*\*\**P* < 0.001). **b**, Correlations between root volume and Euclidean distances of the homozygous F_2:3_ lines from the global average point on the ternary plot. **c**, Comparison of TWAS significance levels between hebQTL-regulated and non-hebQTL-regulated homoeologs. "E-hebQTL" and "N-hebQTL" represent the hebQTLs having and having no genetic effects on root-related traits, respectively. Different letters indicate significant differences (*P* < 0.05) based on least significant difference (LSD) test for multiple comparisons.

Considering that a hebQTL only regulates the homoeolog in the same subgenome, we reasoned that expression variation of the hebQTL-regulated homoeolog might underlie the genetic effects of hebQTLs on root-related traits. To address this hypothesis, we calculated the association significance between the expression variations of the three homoeologs with root-related traits via transcriptome-wide association study (TWAS), respectively (Table S16). The TWAS significance levels of homoeologs regulated by hebQTLs having genetic effects on root-related traits (E-hebQTLs) were significantly higher than those of homoeologs regulated by hebQTLs having no genetic effects (N-hebQTL) (Fig. 5c). In contrast, the TWAS significance levels of non-hebQTL-regulated homoeologs were comparable between triads with E-hebQTLs vs. N-hebQTLs. These results suggest that the genetic effects of hebQTLs on root-related traits could be explained by variation in the expression level of the hebQTL-regulated homoeolog rather than the non-hebQTL-regulated homoeologs.

Notably, the TWAS significance levels of non-hebQTL-regulated homoeologs were significantly higher than those of hebQTL-regulated homoeologs (Fig. 5c), suggesting that hebQTL-regulated homoeologs have weaker effects in regulating root-related traits. This finding is consistent with the lower selection pressure, higher genetic diversity and lower expression levels of hebQTL-regulated homoeologs.

Finally, to further test our results, we selected the triad in which the three homoeologs were annotated as *TaGW2* (*Triticum aestivum Grain Width 2*) to analyse as an example. The expression level of *TaGW2-A1* was lower, while its expression variation was higher than those of its homoeologs *TaGW2-B1* and *TaGW2-D1* (Fig. 6a). We identified four hebQTLs located in subgenome A as regulating the HEB of this triad (Fig. 6b,c). These hebQTLs only had genetic effects on the expression variation of *TaGW2-A1* (Fig. 6d). We selected one of these hebQTLs and validated its genetic effect on HEB using a new F_2:3_ segregation population. Only the relative expression of the hebQTL-regulated homoeolog, *TaGW2-A1*, but not those of *TaGW2-B1* and *TaGW2-D1*, was significantly different between homozygous lines harbouring an alternative allele of this hebQTL (Fig. 6e). The TWAS significance levels of *TaGW2-A1* were much lower than those of *TaGW2-B1* and *TaGW2-D1*, pointing to a weaker effect of *TaGW2-A1* (Fig. 6f). To experimentally validate this conclusion, we generated mutants of the three homoeologs and measured their root-related traits, confirming the weaker function of *TaGW2-A1* in root development compared to *TaGW2-B1* and *TaGW2-D1* (Fig. 6h,i).

**Figure 6.**
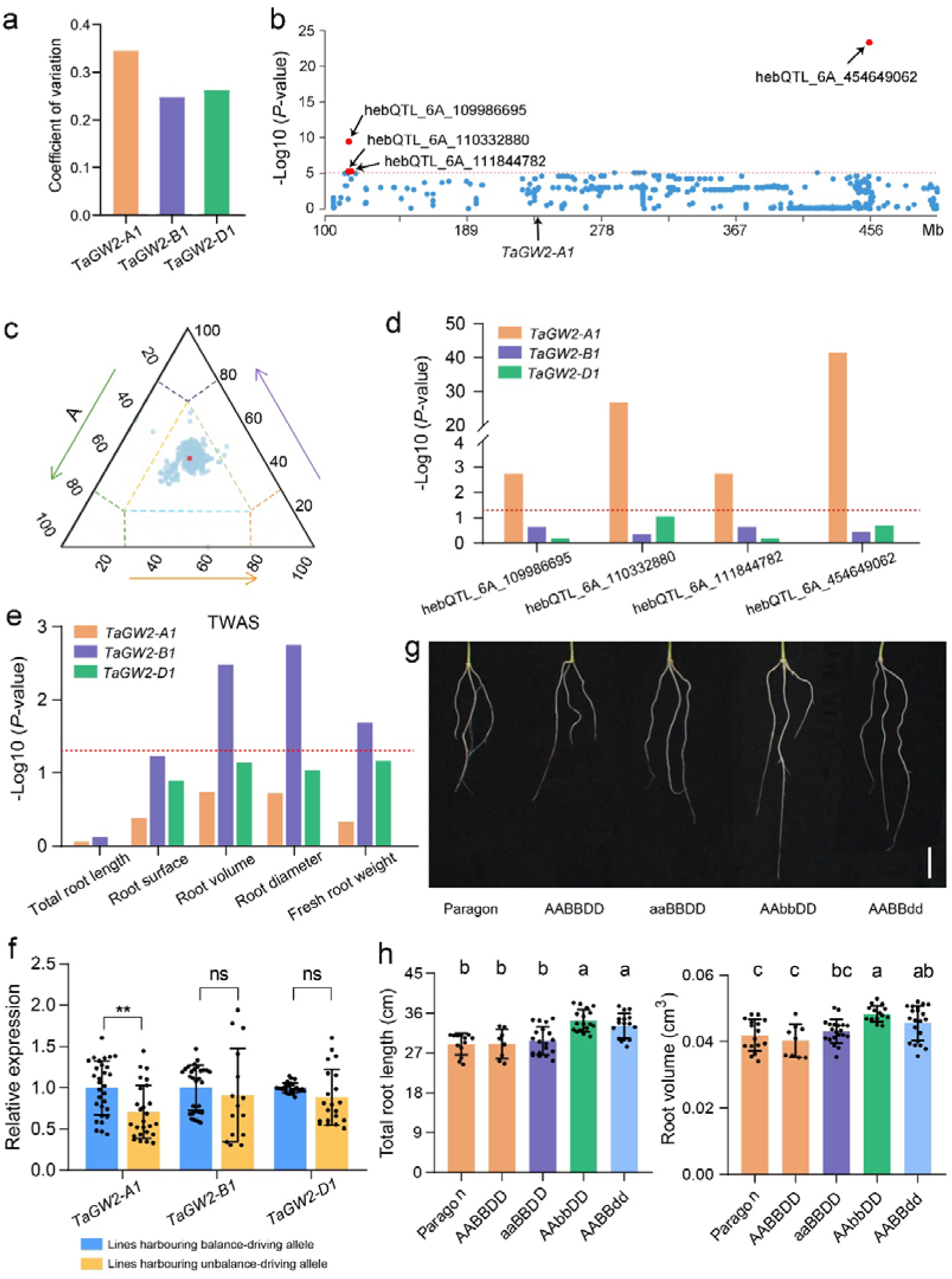
Analysis of *TaGW2* (*Grain Width 2*) as an example. **a**, Comparison of the expression levels of the three homoeologs of *TaGW2* among 406 accessions. **b**, Distribution of the 406 accessions on the ternary plot determined based on the relative expression abundance of the three *TaGW2* homoeologs. Red dot, global average point. **c**, Manhattan plot showing the hepQTLs demonstrated to regulate HEB variation of the *TaGW2* triad. **d**, *P*-values of the genetic effects of the four identified hebQTLs on expression variation of the three *TaGW2* homoeologs. The genetic effects were estimated using the general linear model. **e**, TWAS significance levels for associating expression variation of the three *TaGW2* homoeologs with variations in root-related traits. **f**, Comparison of the expression levels of the three *TaGW2* homoeologs between homozygous F_2:3_ lines harbouring alternative alleles of the most significant hebQTL. \*\**P* < 0.01, ns, *P* > 0.05, determined using Student’s *t*-test. **g**,**h**, Phenotypes of the root-related traits of mutants of the three *TaGW2* homoeologs. Scale bar, 2 cm. In **h**, the bars show the mean ± SD, and different letters indicate significant differences (*P* < 0.05) based on least significant difference (LSD) test for multiple comparisons.

### Downregulation of homoeolog(s) is compensated for by the upregulation of other homoeolog(s) within a triad

The downregulated expressions of hebQTL-regulated homoeologs promote us to investigate the expression dosages of the three homoeologs: that is, whether the downregulation of the hebQTL-regulated homoeolog is compensated for by the upregulation of other homoeologs within a triad. We compared the absolute expression levels of the non-suppressed-homoeolog(s) in accessions in the unbalanced triad category with those of the same homoeolog(s) in accessions in the balanced triad category using the general linear model that accounted for population structure (Fig 7a). Among the 4,148 pairs examined, in 1,592 pairs (38.38%), non-suppressed-homoeologs showed significantly higher expression in accessions in unbalanced vs. balanced triad category (fold change >1.2 and *P*-value <0.05), suggesting that the downregulation of the suppressed-homoeolog was compensated for by upregulation of the non-suppressed-homoeologs (Fig. 7b and Table S17). The proportion of compensated homoeolog-dominant triads ranged from 79.11% to 84.21%, whereas the proportion of homoeolog-suppressed triads ranged from 30.23% to 42.67% (Fig. 7c). To determine whether the downregulation of the suppressed homoeolog was fully compensated for, we compared the sum of expression levels of the three homoeologs between accessions belonging to the unbalanced vs. balanced triad category. The sum of expression levels in accessions in the unbalanced triad category was equal to or even greater than that in accessions in the balanced triad category, ranging from 94% to 98%, suggesting that the downregulation of suppressed-homoeolog was fully compensated for when compensation occurred (Fig. S10, Table S18).

**Figure 7.**
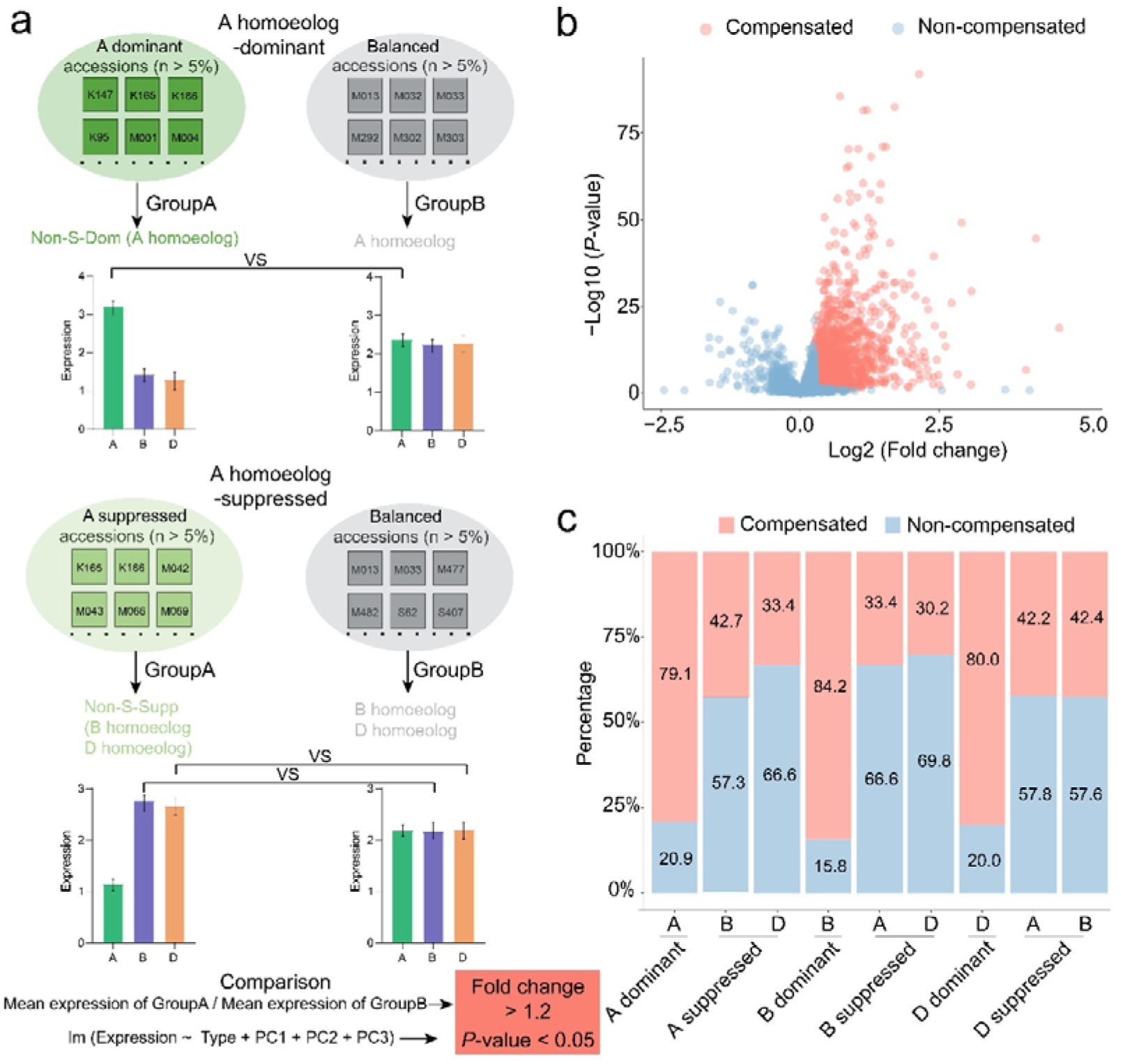
Downregulated expression of hebQTL-regulated homoeolog is compensated for by the upregulation of other homoeologs within a triad. **a**, The model deciphering how to identify the compensation effects, taking an A-homoeolog-dominant triad and an A-homoeolog-suppressed triad as examples. Briefly, for a given triad, the 406 accessions were divided into group A and group B, in which accessions belong to the unbalanced and balanced triad categories, respectively. Then, the expressions of non-suppressed-homoeolog(s) of this triad were compared in the accessions from group A vs group B. **b**, Distribution of the comparison results. "Compensated" represents the comparisons with fold change >1.2 and *P*-value <0.05. **c**, The percentages of compensated and non-compensated triads. The X-axis indicates the homoeologs used to perform comparisons in each of the six unbalanced triad categories.

## Discussion

The differential expression of homoeologs (unbalanced contribution to the total expression level) has been observed in many allopolyploid plants^36–39^ and has been recognized as being advantageous to polyploid plants, as it facilitates adaptive plasticity and fitness. However, how this imbalance is generated is largely known. Specifically, it is currently unclear whether this differential expression is due to a passive process resulting from the accumulation of genetic variants or to an active process resulting from inter-subgenome regulation.

Here, using a newly developed strategy, we identified hebQTLs that regulate variation in HEB among wheat accessions. We determined that a given hebQTL regulates only the expression variation of the homoeolog located in the same subgenome as the hebQTL, which is consistent with the finding that genetic variants in the same subgenome have stronger effects on HEB than those in different subgenomes^33,40^. Notably, the expression levels of hebQTL-regulated homoeologs in accessions harbouring the unbalance-driving allele were lower than those in accessions harbouring the balance-driving allele, suggesting that the downregulated expression of the hebQTL-regulated homoeolog results in HEB. This notion is consistent with our previous finding that suppressed homoeolog expression is the reason for HEB^11^. We determined that non-hebQTL-regulated homoeologs have stronger biological functions, are under higher selection pressure and exhibit lower genetic diversity than hebQTL-regulated homoeologs. Therefore, we suggest that the functions of some hebQTL-regulated homoeologs have been lost or diverged, which resulted in decreased selection pressure on these homoeologs or their regulators and the increased accumulation of genetic mutations, ultimately leading to reduced expression. This notion is consistent with the notion that homoeologs with higher expression within a triad play more important roles in various biological processes. We validated these population-scale findings through functional analysis of *TaGW2*: the A homoeolog is regulated by four hebQTLs and has smaller phenotypic effects than the B and D homoeologs. These results support the notion that suppressed homoeolog expression and the resulting HEB are the results of a passive process attributable to relaxed selection pressure and the accumulation of genetic mutations.

Our identification of hebQTLs suggested that HEB could be inherited and that it might be possible to predict HEB by genotyping the related hebQTLs. Here, the unbalance-driving allele and the downregulated expression of the hebQTL-regulated homoeolog co-segregated in the F_2:3_ population, providing strong evidence for HEB inheritance and the potential for HEB prediction. However, the reports of spatiotemporal specificity of gene expression and differences in HEB categories among wheat tissues appears to partially contradict our finding that HEB is determined by DNA-level variants^11^. We argue that the complexity of the regulation of gene expression underlies this contradiction. For example, the presence of a hebQTL in the gene’s regulating region in the root resulted in its lower expression; however, the genetic regions regulating its expression in other tissues did not accumulate mutations and did not contained a hebQTL, thereby maintaining normal transcription, resulting in different HEB categories among tissues. This hypothesis highlithes the difference in trans-regulatory factors for a given gene’s transcription among tissues. Consistent with this idea, in cotton, the variation in expression bias direction among samples from different timepoints is more likely to be mediated by trans-eQTLs, whereas the stable direction of expression bias is more strongly associated with cis-eQTLs^40^. On the other hand, epigenetic factors that are related to HEB^11,24,31,32^ would also interfere with genotype-based HEB predictions.

In our results, the proportion of hebQTLs having genetic effects on root-related traits is higher than the random controls in subgenomes A and B (Fig. S11a). Notably, 3.86% of these hebQTLs were located in selection sweeps during wheat breeding improvement, and the selected alleles (with higher frequencies in modern cultivars than in landraces) for most of the hebQTLs were unbalance-driving alleles, implying that some HEBs have been exploited in wheat breeding (Fig. S11b,c). The reveal of genetic regulation of HEB provide an avenue to improve wheat breeding by leveraging HEB, e.g. the hebQTL-regulated homoeologs could be upregulated or downregulated by selecting the appropriate alleles of the related hebQTLs. Thereby, HEB reshaping with genetic manipulations become possible.

## Online Methods

### Population transcriptome and identification of genomic variants

The transcriptome data, gene expression values and genomic variants used in this were derived from our previous study and are freely available at the Wheat Genotype and Phenotype Database (WGPD, http://resource.iwheat.net/WGPD/)^34^. Briefly, 406 worldwide bread wheat accessions were collected, seeds of a similar size from each accession were germinated and cultivated on ddH_2_O-soaked filter paper in a germination box, and root-related traits were measured at 14 days after germination in six independent replicates; at least six plants at similar developmental stages were measured per replicate. Six root samples per accession (collected from a single plant per replicate) were combined for bulk transcriptome sequencing. Clean RNA-seq reads were aligned to the bread wheat reference genome (IWGSC RefSeq v1.0) and transcriptome (IWGSC RefSeq v1.1). Using the uniquely mapped reads, gene expression levels were quantified as previously described^11^, and SNPs were called and filtered with the workflow used in our previous study (https://github.com/biozhp/Population_RNA-seq)^34^.

### Definition of homoeolog expression bias categories

The average TPM value of each gene across the 406 accessions was calculated, and the average TPM for the three homoeologs across the triad was normalized to the sum of 1 to acquire relative expression levels as previously described^11^. These values were used to construct ternary diagrams using the R package ggtern (version 3.4.2). The Euclidean distances from the relative transcript abundance of a given triad to each of the seven ideal categories were calculated as previously described^11^ with the rdist function in R (v 4.2.2), and the category with the shortest distance was assigned to this triad. This triad category was called the global category because it was derived from the average TPM value of each homoeolog. The TPM value of each of the three homoeologs in a given accession was calculated as described above; this triad category was called the accession triad category. The global category and the categories of 406 accessions for a given triad were compared to evaluate HEB variation.

### Calculating genetic diversity and estimating selection pressure

To compare the nucleotide diversity and selection pressure surrounding the homoeologs, the nucleotide diversity (π) and *K*_a_/*K*_s_ were calculated using extracted SNPs with MAF >0.05 from publicly available wheat resequencing data^16^ (only hexaploid wheat was used). To calculate nucleotide diversity, the SNPs in the gene region and up- and downstream regions were used, and the π values were determined using VCFtools (v0.1.16)^41^. The sliding window size and step size were set to be equal to the investigated gene length. To calculate *K*_a_/*K*_s_, only the SNPs in the gene region were used, and the missense and synonymous SNPs were annotated using SnpEff (v5.2)^42^; subsequently, the *K*_a_/*K*_s_ values for each gene were calculated using selectionTools (v1.1)^43^.

### Identification of hebQTLs

For a given dynamic triad, the 406 accessions were projected onto the ternary plot based on the relative expression abundance of the three homoeologs. The ternary plot was divided into six regions based on the global average position (Fig. 3e). Two head-to-head regions were considered to be a single GWAS region, generating three GWAS regions for each triad. The accessions in each of the two sub-regions of each GWAS region were counted, and only the GWAS regions containing more than 21 accessions in each of the two sub-regions (5% of the total accessions) and more than 120 accession in the two sub-regionswere maintained for further analysis. The ED values of the accessions located in the sub-region driving for balance were regarded as positive values, while those of the accessions in the opposite sub-region were regarded as negative value. This strategy transforms the extent of imbalance into one-dimensional values for a given triad.

The transformed ED values of each GWAS region for a given triad were associated with the genetic variants identified in our previous study by GWAS. GWAS was performed with the mixed linear model (PCA + kinship) implemented in GEMMA^44^. The PCA and kinship matrix were generated using GAPIT^45^ with default parameters. A Bonferroni-corrected threshold probability based on individual tests was calculated to correct for multiple comparisons using 1/*N*, where *N* is the number of SNPs in the accessions in the relative GWAS-region.

To assign the significantly associated SNPs to hebQTL intervals, a revised two-step method^46^ was employed as follows: (1) all of the associated SNPs were grouped into one cluster if the distance between two consecutive SNPs was <10 kb, and the clusters with at least three significantly associated SNPs were considered to be candidate hebQTLs; (2) a candidate hebQTL in linkage disequilibrium (LD) (*r*^2^ > 0.1) with other more significant candidate hebQTLs were regarded as associations introduced by the LD structure and were removed.

### Estimation of the genetic effects of hebQTLs on gene expression and root-related traits

The genetic effects of hebQTLs were estimated using our previously described model, which was developed to estimate the genetic effects of a given LD block on the investigated root-related traits^35^. Briefly, the general linear model, in which the population structure was included as a fixed effect, was calculated with the lmerTest (v3.1.3) package in R (v 4.2.2) using the following equation:

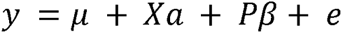

In this equation, *μ* represents the overall mean, *Χ* represents the alleles of the investigated hebQTL and *P* represents the top three principal components. *Xα* and *Pβ* were used as fixed effects. When estimating the genetic effect of a hebQTL on gene expression, *y* represents the TPM value of the gene; when estimating the effect on root-related traits, *y* represents the measured trait values. Notably, the root-related trait values were normalized with the general linear model to exclude the effects of kernel weight. To improve the accuracy, the accessions harbouring alternative alleles were counted, and only the hebQTLs where the two numbers were both larger than 30 were maintained.

### Validation of hebQTLs using segregation populations

Two F_2:3_ bi-parental segregation populations were constructed by crossing Lianmai2 with Safha3 (to validate Triads401, Triads10304 and Triads10311) and Qingfeng1 with Yannong22 (to validate *TaGW2*). The growth conditions and root-related traits were measured using the same procedures employed to phenotype the 406 accessions. Whole roots of individuals were sampled and frozen in liquid nitrogen. Total RNA was isolated from the samples using RNAiso Plus (Takara, Cat. #9108) and reverse transcribed into cDNA with random primers using Evo M-MLV reverse transcriptase (Accurate Biology, Cat. #AG11728). qRT-PCR was performed to measure the expression level of the target gene using TB Green Premix Ex Taq II (Takara, Cat. #RR820A). The primer sequences are listed in Table S12. For genotyping, Kompetitive Allele Specific PCR (KASP) markers (Table S13) were developed for each hebQTL using PolyMarker (http://www.polymarker.info/) and used to assign the genotype of each individual in the F_2:3_ populations. The relative expression and phenotypic values of the individuals carrying alternative alleles were compared using the Student’s *t*-test; only homozygous lines were selected for this analysis.

### Association analysis between Euclidean distance values and root-related traits

Only regions used in GWAS were used for association analysis. The transferred ED values of accessions in each GWAS region for a given triad were associated with root-related traits. Association analysis was performed with the general linear model and the equation used to estimate genetic effect described above, in which *Χ* represents the transferred ED values and *y* represents the investigated trait.

### Transcriptome-wide association study (TWAS)

TWAS, a technique to examine the association between gene expression variation and phenotype variation among a population, is usually performed with software developed for GWAS in plants by transforming the continuous variable (gene expression levels) into a discrete variable ("0" and "1"). However, this transformation results in the loss of some gene expression information. Here, the linear mixed model, the fundamental model in GWAS, was used to exploit the continuous gene expression variations and to adjust for population structure and family relatedness with the lmerTest (v3.1.3) package in R (v 4.2.2) as follows:

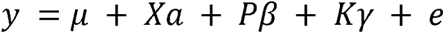

In this equation, *y* represents the values of the investigated phenotype, *μ* represents the overall mean, *Χ* is a continuous variable that represents gene expression values, and *P* and *K* are the PCA and relative kinship matrix generated with GAPIT^45^ using the parameters “PCA.total=3”. The top three principal components were used to build the *P* matrix for population-structure correction. The *K* matrix was used to correct the potential familial relatedness. *Xα* and *Pβ* represents fixed effects, and Κγ represents random effects. The root-related trait values were normalized with the general linear model to exclude the effects of kernel weight. The genes with false discovery rate (FDR)-adjusted *P-*value <0.01 were considered to be candidates significantly associated with the investigated phenotypes. Only genes with TPM > 0.5 in >95% of accessions were used for TWAS.

### Phenotypic evaluation of *TaGW2* mutants

Wild-type (AABBDD) progeny and single *TaGW2* mutants (aaBBDD, AAbbDD and AABBdd) were derived from BC_4_ near-isogenic lines produced by crossing and back-crossing of the TILLING mutant lines (Kronos2335 for *TaGW2-A1*, Kronos0341 for *TaGW2-B1*, Cadenza1441 for *TaGW2-D1*) to the cultivar Paragon as described previously^47^. Seeds of a similar size from each mutant were germinated and cultured on ddH_2_O-soaked filter paper for three days. The seedlings were transferred to containers (40 × 30 × 12 cm) filled with 1/2× Hoagland solution with punched lightproof plastic covers and cultivated under 16 h light (2000 Lux)/8 h dark cycle at 22℃/16℃ day/night (50% relative humidity). The Hoagland solution was changed weekly. The total root length, root surface, root volume and root diameter were examined with a Wseen LA-S image system (Hangzhou Wseen Testing Technology Co. Ltd.), and primary root length was measured manually.

### Analysis of expression dosage compensation

To investigate whether the downregulation of the suppressed homoeolog for a given unbalanced triad is compensated for by the upregulation of other homoeologs within the triad, the 406 accessions were divided into two groups, group A and group B, comprising accessions harbouring unbalanced and balanced triad category, respectively (Fig. 7a). Only the triads where the accession numbers in groups A and B are both more than 21 (5% of the total accessions) were maintained. The TPM values of the non-suppressed homoeolog in the accessions from group A were compared with those of the same homoeolog in the accessions from group B using the general linear model with the lmerTest (v3.1.3) package in R (v 4.2.2) as previously described^48^, in which the population structure was included as a fixed effect with the following equation:

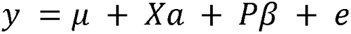

In this equation, *y* represents the TPM value of the investigated homoeolog, *μ* represents the overall mean, *Χ* represents the category (group A or B) and *P* represents the top three principal components generated with GAPIT (v3.1). *Xα* and *Pβ* were used as fixed effects. The sum of expression levels of the three homoeologs across the triad were compared between group A and group B accessions with the same model to investigate whether the downregulation of the suppressed homoeolog was fully compensated for.

### Identification of homoeologous genes among the three subgenomes and GO enrichment analysis

The 1-to-1 homologous gene table among three subgenomes of Chinese Spring was downloaded from Triticeae-GeneTribe (http://wheat.cau.edu.cn/TGT/download.html). The homology inference among genetically similar genomes was calculated by integrating the collinear block scores, coding sequence similarity and annotation quality information^49^.

GO enrichment analysis was performed with the "GOEnrichment" module of Triticeae-GeneTribe (http://wheat.cau.edu.cn/TGT/)

## Acknowledgements

We thank Cristobal Uauy for his insightful comments and suggestions on the work. The research was funded by grants from the China Postdoctoral Science Foundation (2021T140566). This study was supported by the High-Performance Computing of NWAFU.

## Author contributions

X.W. designed the study. X.W., S.X. and W.J. collected the data. Y.L., P.Z. and X.W. analysed the data. J.S. generated the mutants of *TaGW2*. W.H. and M.C. generated the F_2:3_ populations, collected the phenotypes of F_2:3_ populations and *TaGW2* mutants, and performed the qRT-PCR in the F_2:3_ populations. P.B. and X.S. provided intellectual input. X.W. wrote the initial draft of the manuscript. All authors reviewed the manuscript and gave approval for submission of the final version.

