## Supplemental Figures for "Intra-subgenome regulation induces unbalanced expression and function among bread wheat homoeologs"

**Supplemental Figures**
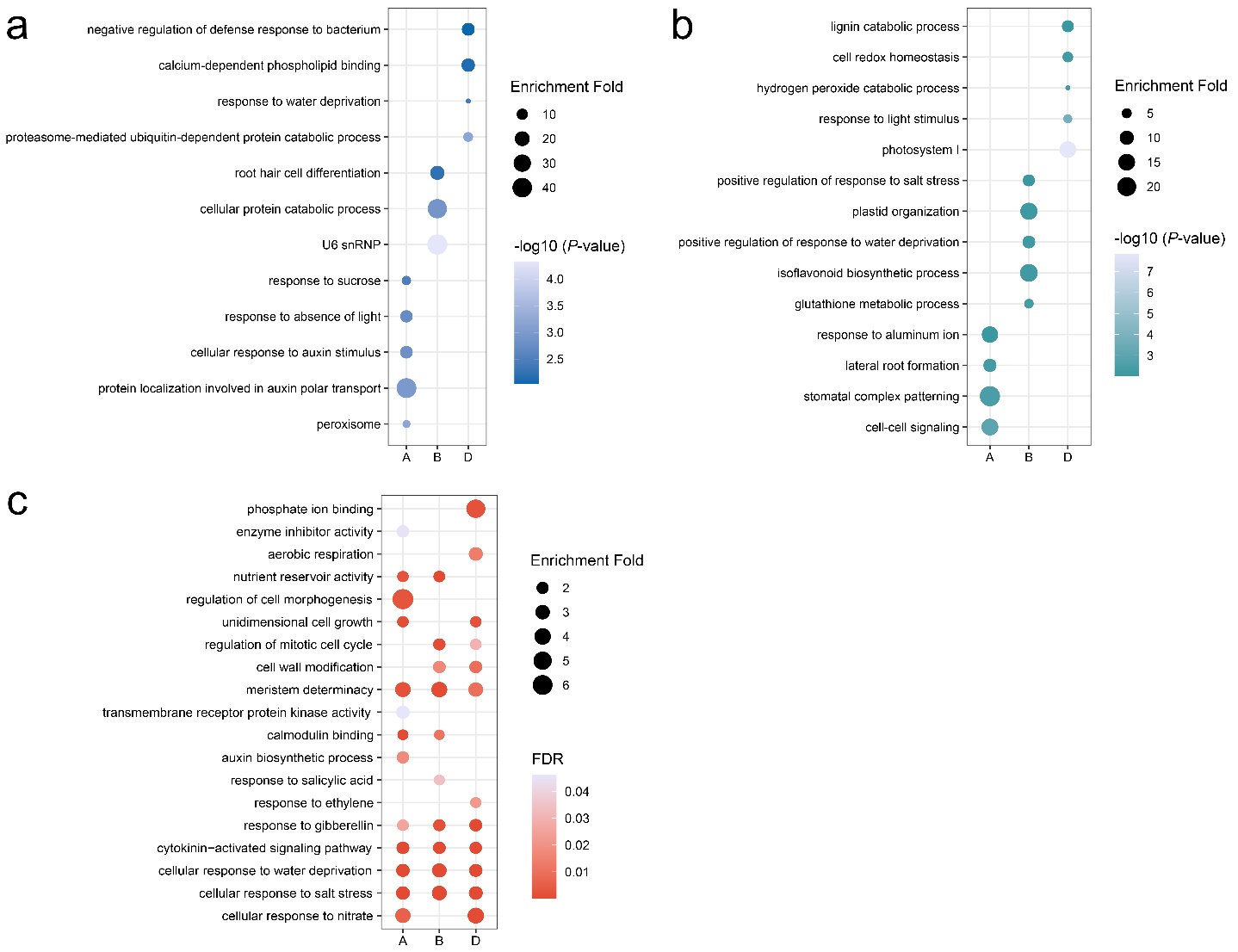

Figure S1. Gene ontology enrichment of the "private" genes. **a-b**, GO enrichment of the subgenome-private genes in Fig. 1c (**a**) and Fig. 1e (**b**). The full list of significantly enriched GO terms is included in Table S2 and Table S4. **c**, GO enrichment of the "private" genes in Fig. 1f. The full list of significantly enriched GO terms is included in Table S6.

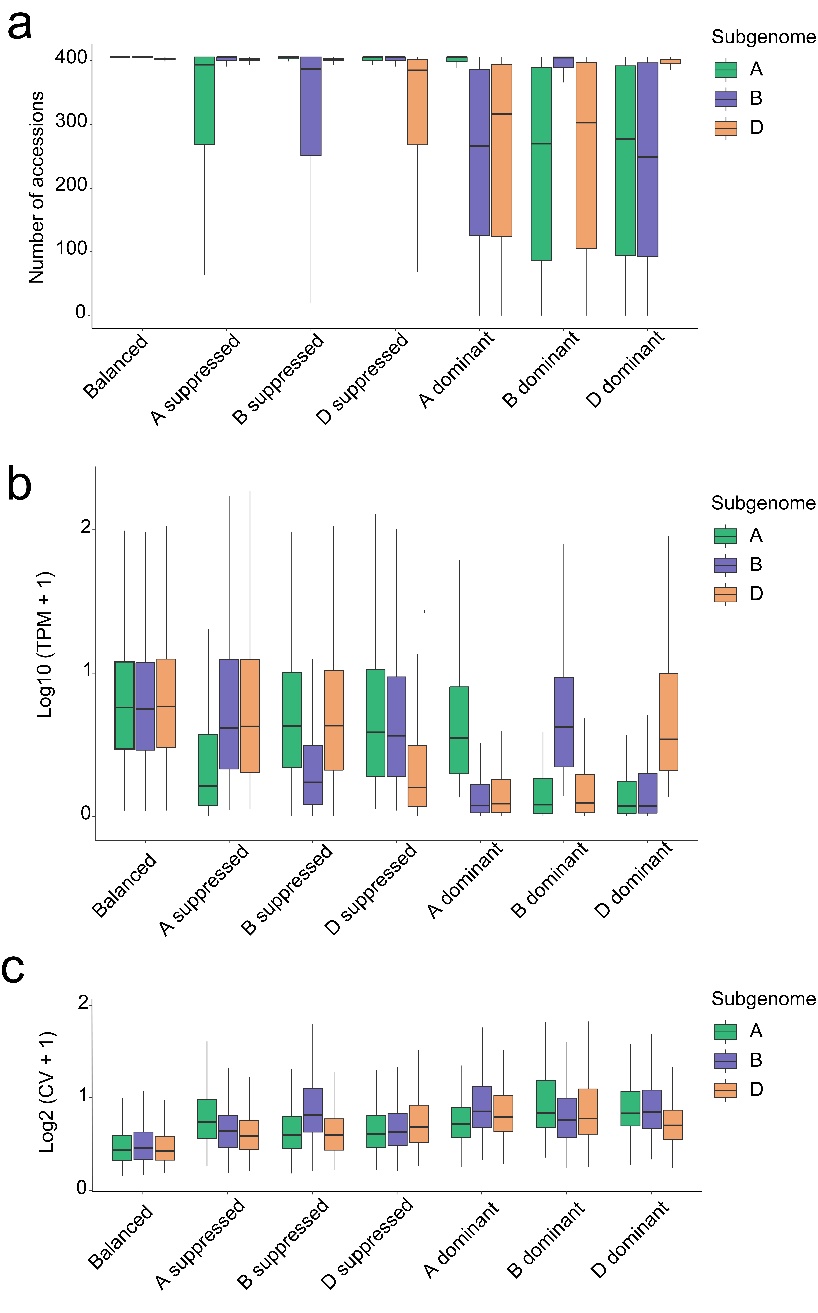

Figure S2. Expression status of the three homoeologs from seven triad categories. **a**, Expression range of the three homoeologs, i.e., the number of accessions in which they are expressed. **b**, Absolute expression levels of the three homoeologs. **c**, Coefficient of variation (CV) of gene expression levels among the 406 accessions.

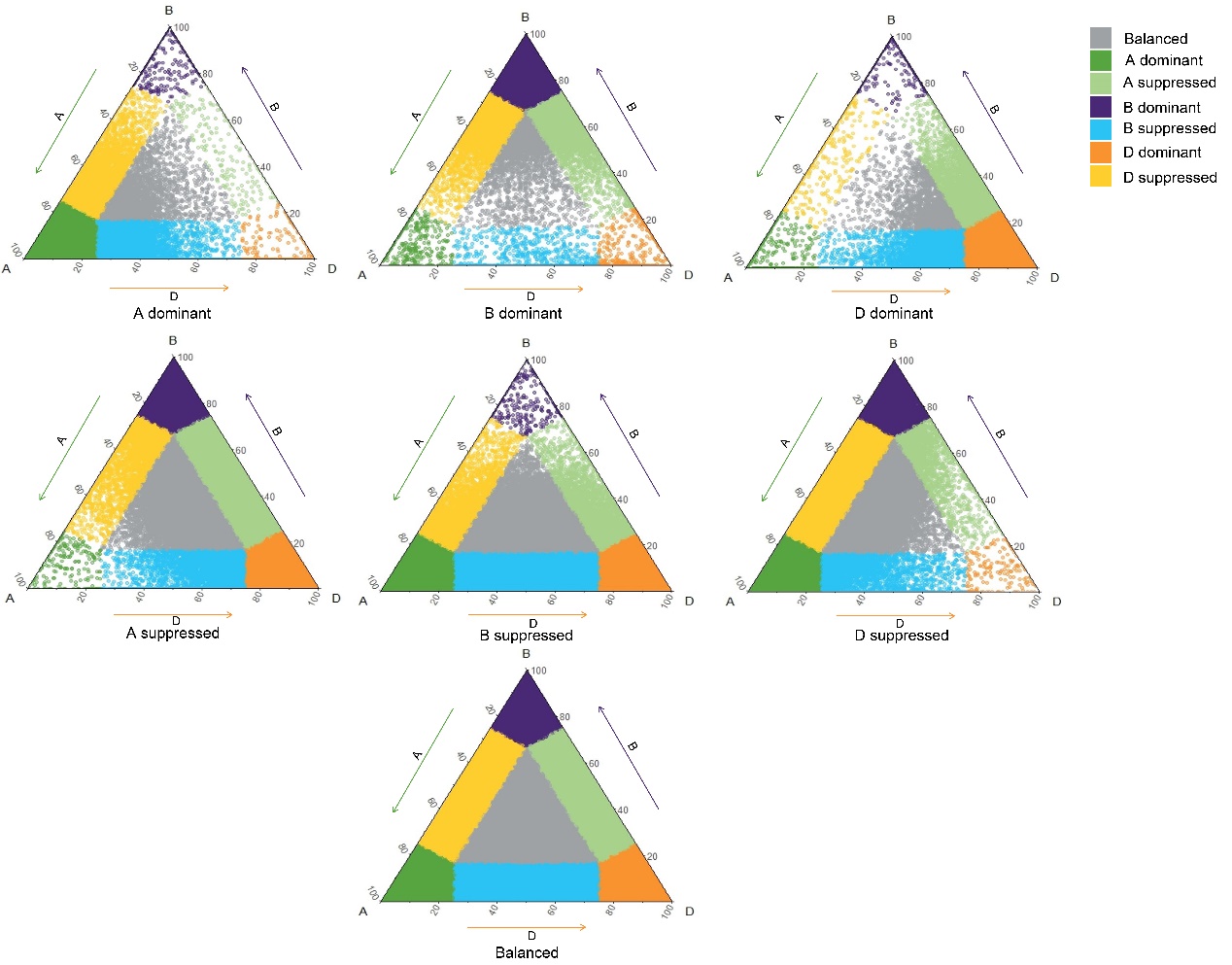

Figure S3. Variation in homoeolog expression patterns among the 406 accessions. The seven ternary plots represent the seven triad categories determined based on the mean expression values of the three homoeologs among the 406 wheat accessions. For each triad in a given global category, the relative expression abundance of the three homoeologs was calculated based on their absolute expression levels in each accession and projected onto the ternary plot. Therefore, each triad produced 406 points on the ternary plot.

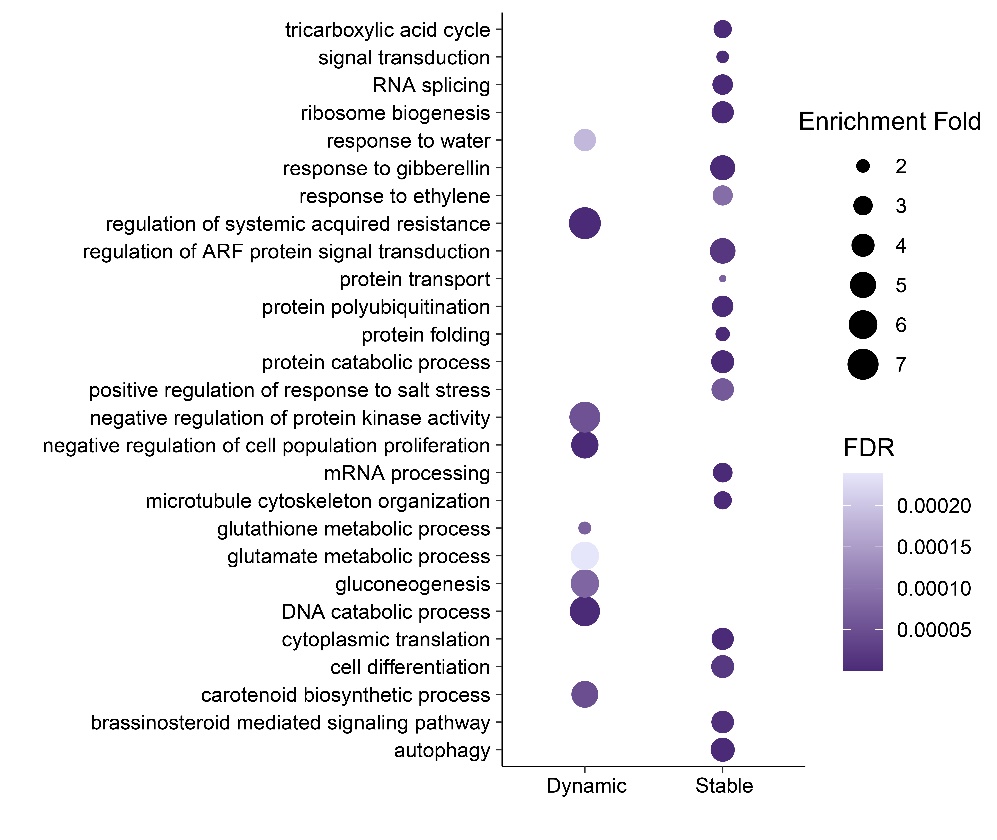

Figure S4. Gene Ontology enrichment analysis of the genes from dynamic and stable triads. The full list of the significantly enriched GO terms is included in Table S10.

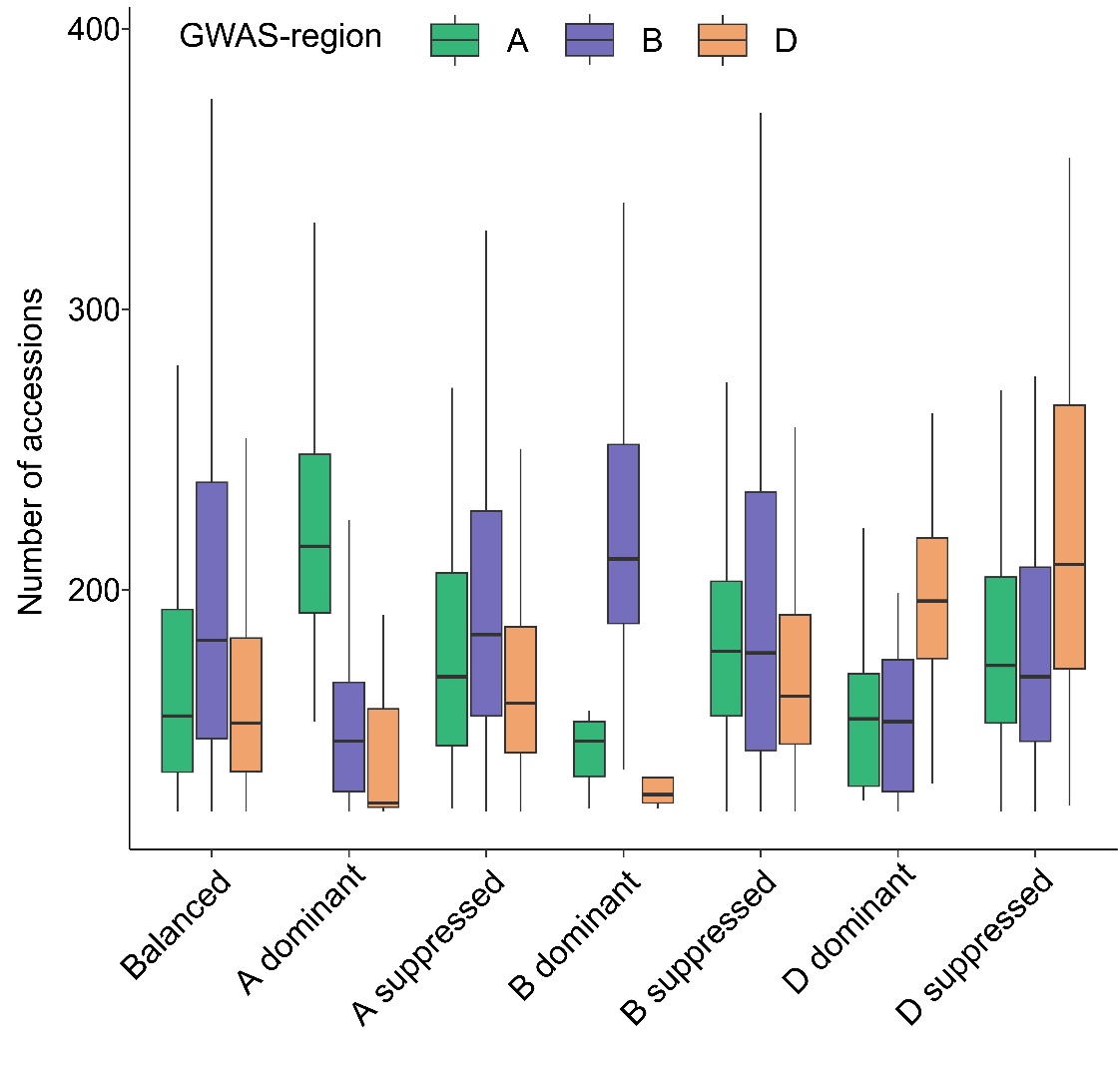

Figure S5. Number of accessions in each of the three GWAS regions. A, B and D represent the GWAS region containing the vertex A, B and D of the ternary plot in Fig. 3**e**, respectively.

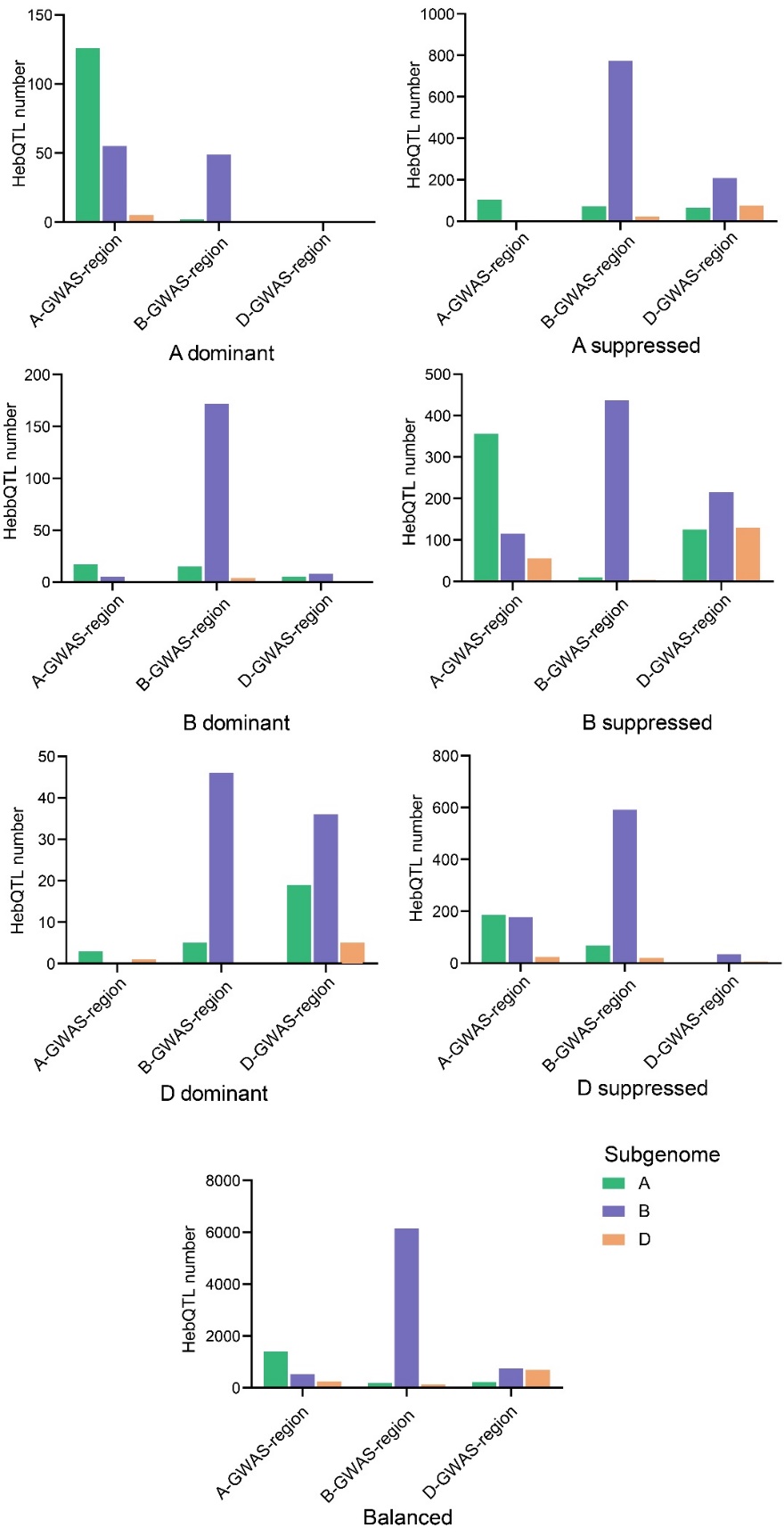

Figure S6. Number of identified hepQTLs in each of the three GWAS regions for the seven triad categories. The A-, B- and D-GWAS regions represent the GWAS region containing vertices A, B and D of the ternary plot in Fig. 3e, respectively. Subgenomes A, B and D indicate which subgenome the identified hepQTLs located on.

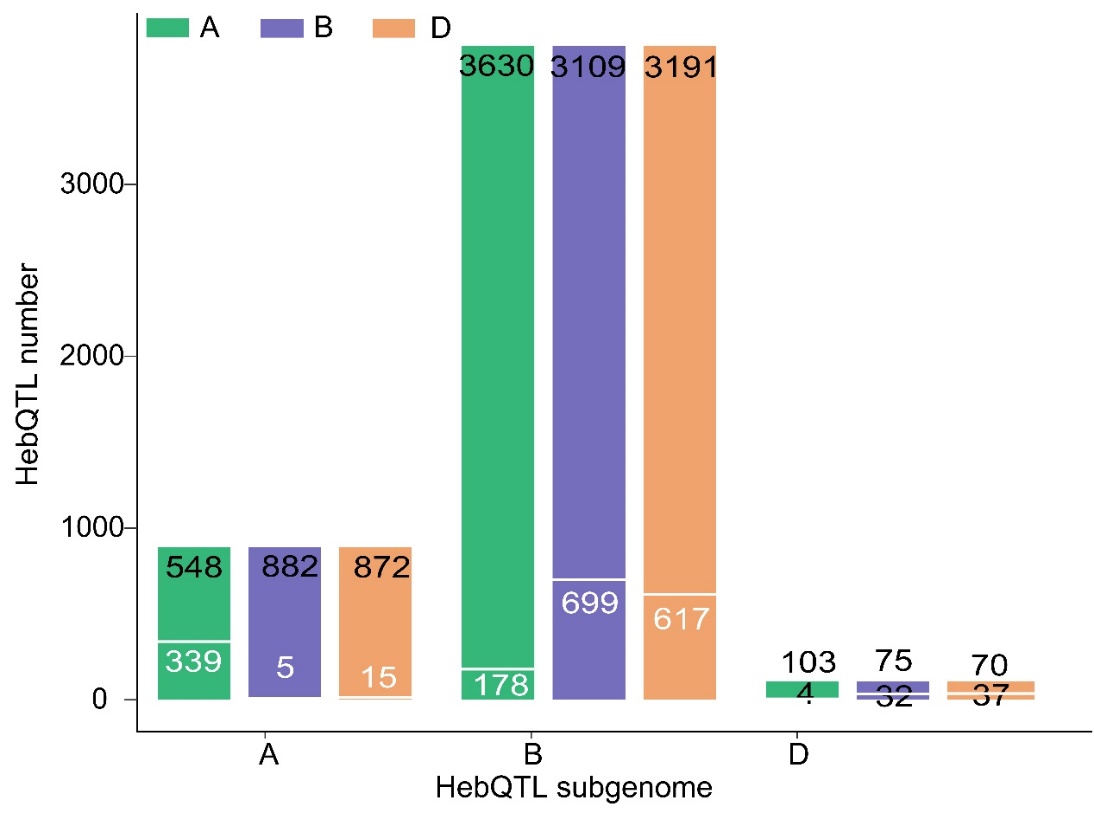

Figure S7. Overlap between the triad's hepQTLs and the eQTLs of the three homoeologs. Black and white numbers indicate the number of overlapping and non-overlapping hepQTLs, respectively.

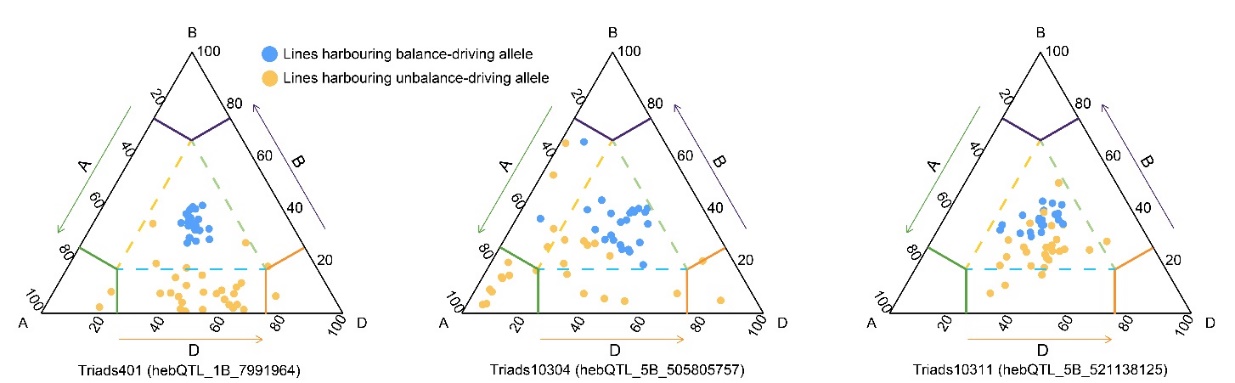

Figure S8. Distribution of homozygous F_2:3_ lines carrying alternative hepQTL alleles on the ternary plot. The position of each line was determined based on the relative expression levels of the three homoeologs, as detected by qRT-PCR.

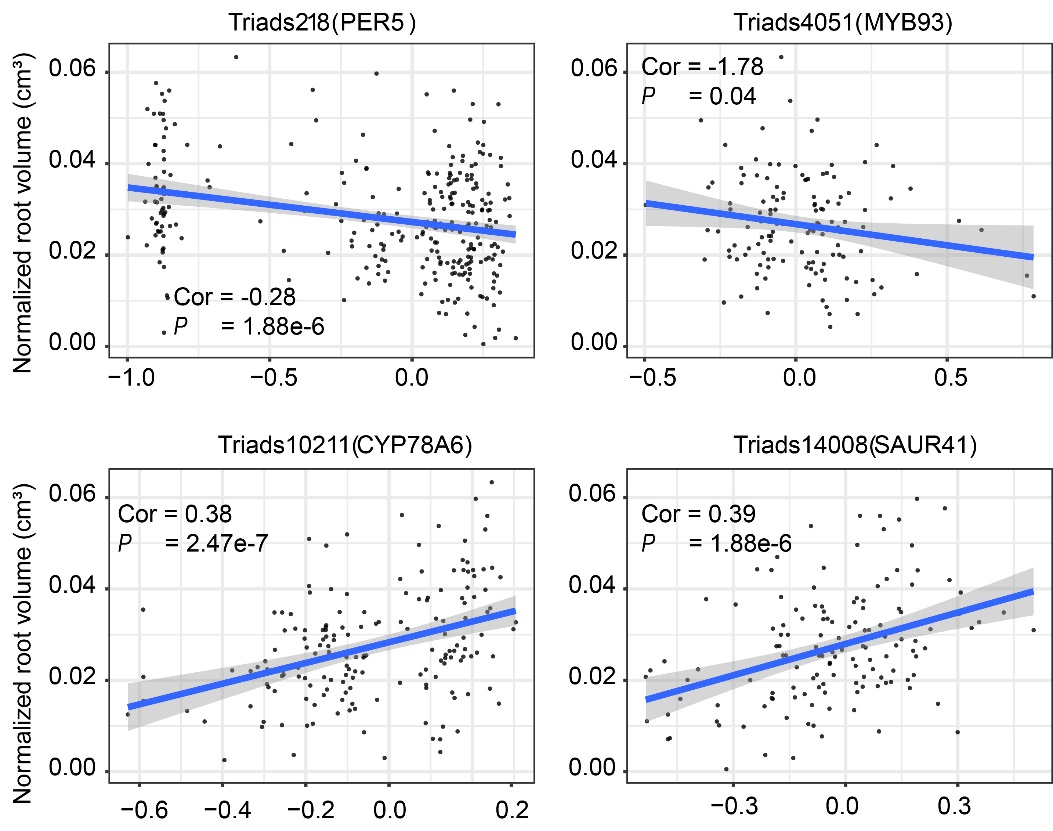

Figure S9. Association between Euclidean distance values and root-related traits in four triads. For a given F_2:3_ line, the Euclidean distance between the triad’s position of this line and its global average position on the ternary plot was calculated. The gene names represent the wheat homoeologs of *AtPER5*, *AtMYB93*, *AtCYP78A6* and *AtSAUR41*.

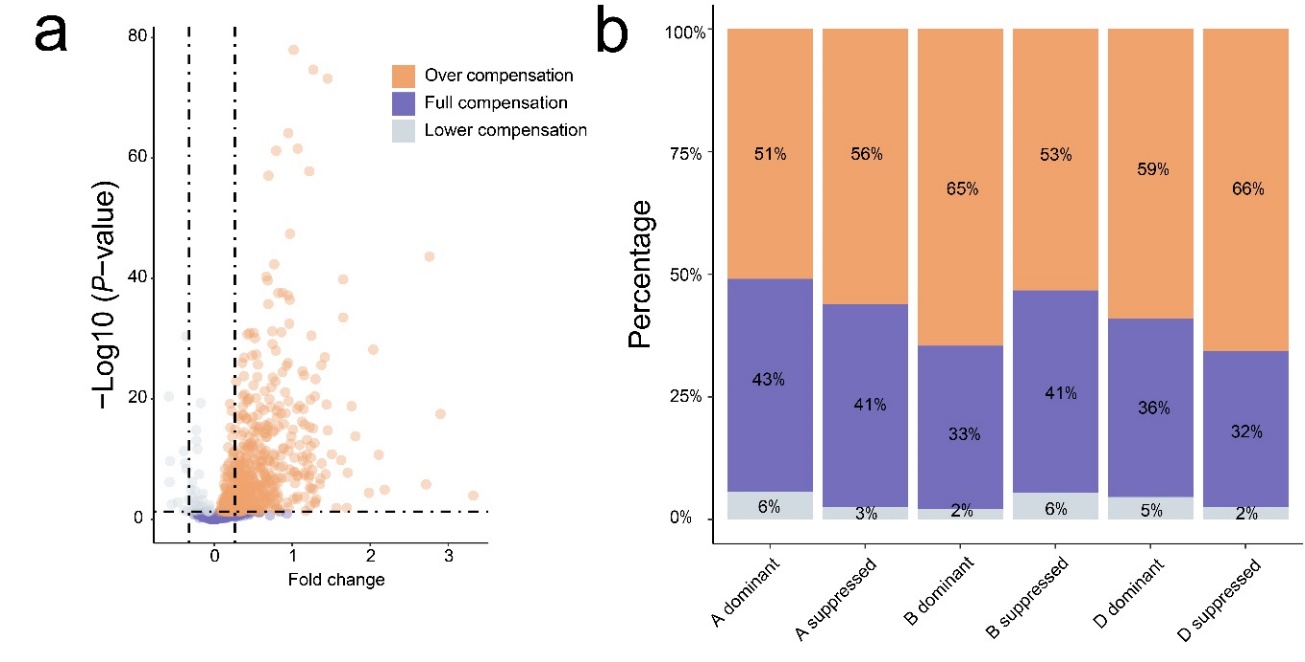

Figure S10. The extent to which the downregulation of the suppressed-homoeolog is compensated for by the upregulation of other homoeologs within a triad. Only triads in which the downregulation of the suppressed-homoeolog was compensated for were used. **a**, Comparisons of the sum expression levels of the three homoeologs between accessions in unbalanced vs. balanced category. "Over compensation", "Full compensation" and "Lower compensation" indicates that the sum expression levels in accessions in the unbalanced category were higher than, comparable to and lower than those in accessions in the balanced category, respectively. The comparisons were performed with the general linear model. **b**, Proportions of "Over compensation", "Full compensation" and "Lower compensation" triads.

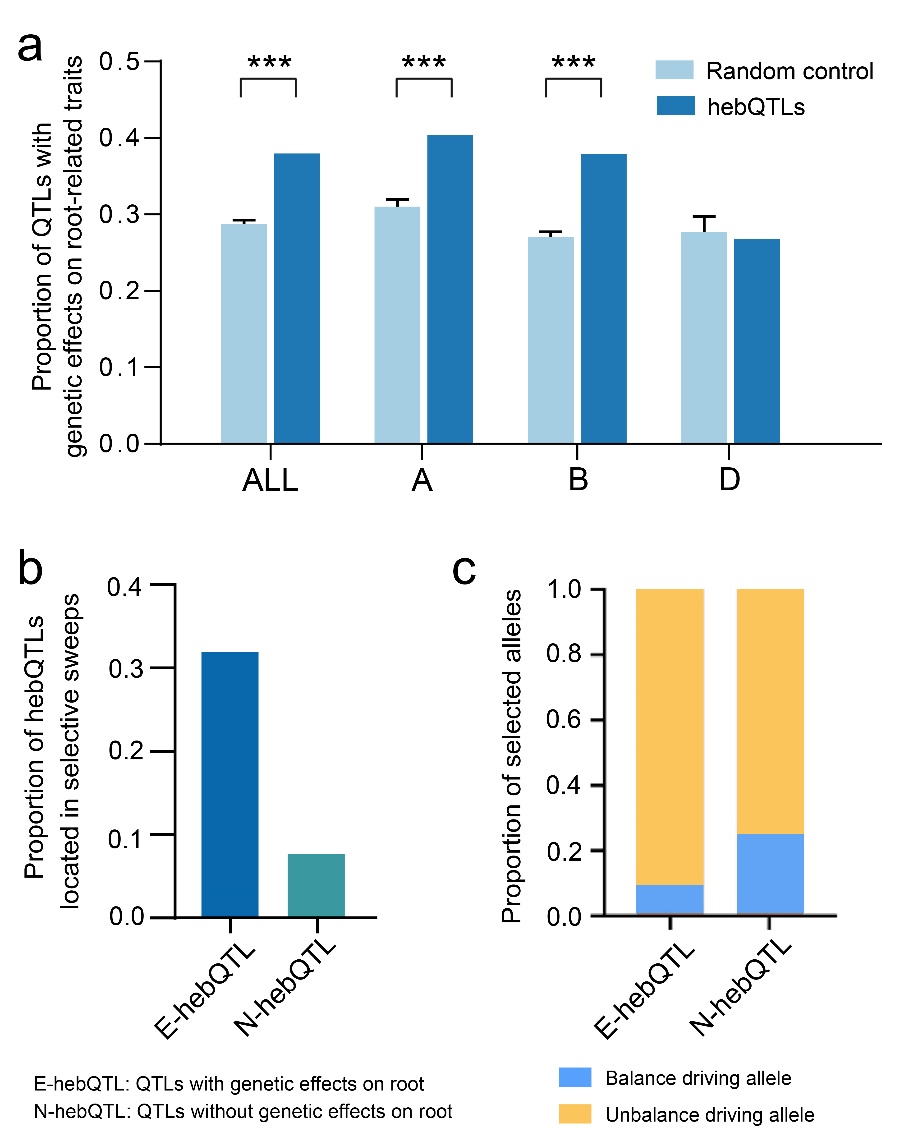

Figure S11. The hepQTLs with genetic effects on root development were involved in breeding selection. **a,** The proportion of hepQTLs with genetic effects on root-related traits. The random control was generated with 1,000 permutation tests by sampling an equal number of genomic linkage blocks. **b,** The proportion of hepQTLs located in selective sweeps. **c,** The proportion of the allele selected by wheat breeding.
